# Bacterivorous nematodes correlate with soil fertility and improved crop production in an organic minimum tillage system

**DOI:** 10.1101/2020.06.08.139840

**Authors:** Jan H. Schmidt, Johannes Hallmann, Maria R. Finckh

**Affiliations:** Ecological Plant Protection Group, University of Kassel, Witzenhausen, Germany; Institute for Epidemiology and Pathogen Diagnostics, Julius Kühn-Institute, Federal Research Centre for Cultivated Plants, Braunschweig, Germany

## Abstract

Organic farming systems are generally based on intensive soil tillage for seed bed preparation and weed control, which in the long-term often leads to reduced soil fertility. To avoid this, organic farming systems need to adopt conservation agriculture practices, such as minimum tillage and diligent crop rotations. However, minimum tillage generally delays soil warming in spring causing reduced nitrogen mineralization and thus poor plant growth. This negative effect needs to be compensated. We hypothesize that, in a diverse crop rotation, organic minimum tillage based on frequent cover cropping and application of dead mulch will improve soil fertility and thus crop production as confirmed by a number of chemical and biological soil indicators.

We made use of two long-term field experiments that compare typical organic plough-based systems (25 cm) with minimum tillage systems (<15 cm) including application of transfer mulch to potatoes. Both tillage systems were either fertilized with compost or equivalent amounts of mineral potassium and phosphate. In 2019, soil samples from both fields were collected and analyzed for soil pH, organic carbon, macro-, micronutrients, microbial biomass, microbial activity and total nematode abundance. In addition, performance of pea in the same soils was determined under greenhouse conditions.

Results from the field experiments showed an increase of macronutrients (+52%), micronutrients (+11%), microbial biomass (+51%), microbial activity (+86%), and bacterivorous nematodes (+112%) in minimum tillage compared with the plough-based system. In the accompanying greenhouse bioassay, pea biomass was 45% higher under minimum than under plough tillage. In conclusion, the study showed that under organic conditions, soil fertility can be improved in minimum tillage systems by intensive cover cropping and application of dead mulch to levels higher than in a plough-based system. Furthermore, the abundance of bacterivorous nematodes can be used as a reliable indicator for the soil fertility status.

## Introduction

Organic farming systems are generally based on intensive soil tillage for seed bed preparation and weed control, which in the long run often leads to reduced soil fertility [1]. Although intensive soil tillage increases microbial turnover rates and thus nutrient availability required for plant growth, long-term intensive soil tillage can cause depletion of the soil organic carbon content and thus reduced soil fertility [2]. For a long-term improvement of soil fertility and its maintenance at a sustainable level, organic production systems need to reduce the frequency and intensity of soil tillage and increase the organic matter supply to the soil. The resulting accumulation of organic carbon will likely increase the microbial activity and thus result in accelerated nutrient cycles [3–5]. However, minimum tillage generally tends to delay soil warming in spring, and therefore N-mineralization rates are often too low to meet the demand of the crops, especially in temperate climates [6]. That’s why applying conservation agriculture methods, i.e. the simultaneous application of minimum tillage, crop rotations, and residue retention, to organic farming may not necessarily improve soil fertility, even after 10 years of adaptation to the system [7]. Similary, Krauss et al. [5] reported that yield of winter wheat, silage maize, and spelt in an organic long-term experiment was still 10% lower even more than 10 years after transition to reduced tillage compared to standard moldboard ploughing, even though manure compost and slurry had been frequently applied Although nutrient levels and biological soil components were generally higher under reduced tillage compared to plough tillage in the top 10 cm soil, the massively enhanced weed competition under reduced tillage likely reduced crop yields. Thus, organic minimum tillage systems need to be modified in order to provide sufficient levels of nutrients and weed control at the same time [8].

Two options to achieve this might be the use of legume and non-legume cover crops and mulches. Cover crops are known among others to conserve the nutrients of the previous crop for the following crop, increase the organic matter content, stimulate microbial activity and suppress weeds [1]. Especially leguminous cover crops and cover crop mixtures of brassicas with legumes have shown positive effects on microbial biomass and activities as well as specific enzyme activities independent of the climatic region and weather conditions [9]. Furthermore, the use of cover crops can reduce weed seed banks in minimum tillage systems similar to levels in plough systems [10]. Organic mulch applications, referred here as the harvest of cover crops and their subsequent application to a specific crop or field, have been shown to contribute substantially to soil fertility in organic minimum tillage systems [6]. All those measures also protect the soil from a range of environmental impacts, such as drought, wind and water erosion or even plant diseases [11].

In combination with a long-term organic fertilizer strategy, such cropping systems should result in more sustainable cropping systems in which nutrient cycles are almost closed. For example, application of high quality and certified composts that are free of pathogens, weeds, and toxic compounds can contribute to a better plant performance in minimum tillage systems. Besides nutrients, composts introduce additional microorganisms to the systems that may contribute to the suppression of soil-borne diseases and should therefore enhance the overall soil fertility [12]. However, the evidence of disease suppression and the resulting soil fertility improvement through the use of composts often failed under field conditions in temperate climates due to variable environmental conditions and inadequate application rates of composts [13,14]. Thus, long-term field trials are required for a deeper understanding of the importance of compost in disease suppression and soil fertility improvement [15].

Soil fertility, which in this context is used synonymous for soil quality and soil health, can be assessed through chemical and biological indicators, such as organic carbon, pH, micro- and macronutrients, microbial biomass, or microbial respiration [16]. Furthermore, free-living nematodes are considered important indicators of soil quality [17–20]. Different feeding types of nematodes occupy different niches within the soil food web and hence, their classification and enumeration can determine certain carbon pathways. In a recent review, Bünemann et al. [16] pointed out that biological indicators are rarely used to assess soil health and quality and that most of the commonly used indicators are “black box” indicators, such as C_mic_ and microbial respiration. They further criticize that such assessments are rarely linked to specific ecosystem services, which impedes the evaluation of their suitability as soil quality and health indicators.

Here we investigated two long-term experiments that were set up in adjacent fields in 2010 and 2011 to assess the effects of an organic minimum tillage system on chemical and biological soil properties over time. The study specifically addressed the question, whether a crop rotation that includes cover crops and mulch applications can maintain or even improve soil fertility and if this can even be further improved by the regular application of compost. Furthermore, the study investigated which chemical and biological parameters were best linked with biomass production in a pea (*Pisum sativum* L.) bioassay and therefore could serve as indicator for soil fertility. The study compared a typical plough-based system (25 cm) with a minimum tillage system (max. 15 cm), whereas the minimum tillage system comprised applications of transferred dead mulch to potatoes (experiment 1: 2014, 2018; experiment 2: 2015, 2019) (Fig 1). The second factor analyzed was the application of yard waste compost at a rate of ∼5 t (ha a)^−1^ dry matter (DM) compared to equivalent amounts of mineral phosphorus (rock phosphate) and potassium (K_2_SO_4_) fertilization. Soils of both field experiments were evaluated in 2019. This was year 9 of experiment 1 with clover-grass as main crop and year 8 of experiment 2 with potatoes as main crop. The soil of both field experiments was further analyzed in the greenhouse for biomass production in a pea bioassay. We hypothesized that i) minimum tillage with mulch to potatoes increases soil fertility compared with a plough-based inversion tillage system without mulch, ii) regular compost application improves chemical and biological soil parameters compared to mineral fertilization, and iii) soil fertility indicators, including bacterivorous nematodes, are positively correlated with pea biomass production and reduced root disease severity in the greenhouse bioassay.

**Fig 1.**
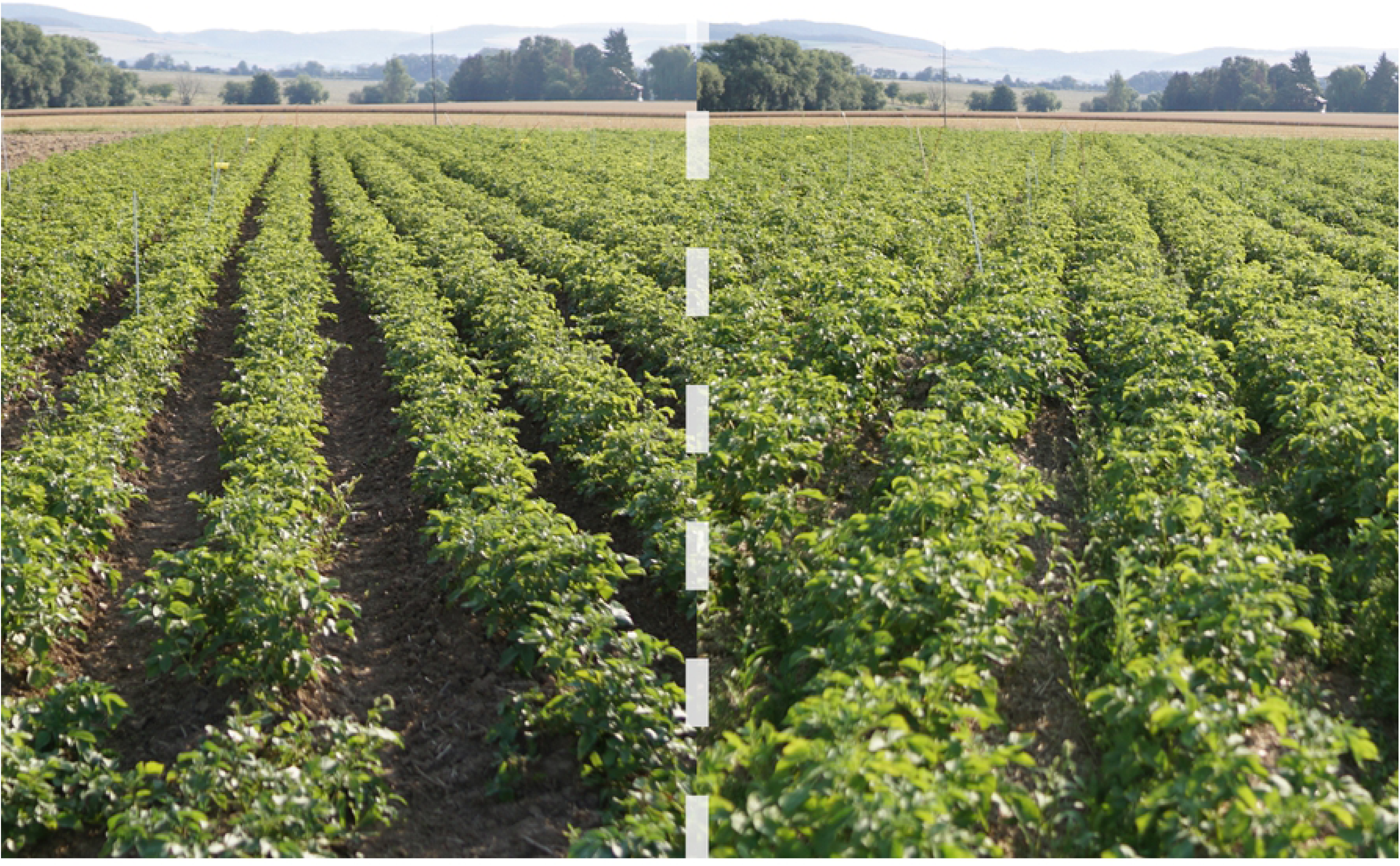
Soil fertility indicated by potato canopy closure on July 24th 2019 prior to flowering (BBCH 59). Left: Plough tillage without mulch; right: Minimum tillage with mulch. Potatoes were planted end of May. Foto: S. Junge

## Materials and Methods

The two long-term experiments were originally started in 2010 and in 2011 in adjacent fields located on the organic experimental farm of the University of Kassel in Neu-Eichenberg (51°22’51”N, 9°54’44”E, 231 m ASL with an eastern incline of 3%). The soil type is a Haplic Luvisol with 3.3% sand, 83.4% silt and 13.3% clay (USDA classification Zc). Liming (CaCO_3_) was applied to all treatments at 2 t ha^−1^ in August/September 2019.

The experiments have been described in detail in Schmidt et al. [21] and all crops grown since the start are shown in Table 1. In brief, both experiments consist of a split-plot design with four replicates and tillage as main factor (12 × 60 m^2^): 1) minimum tillage by chisel ploughing or shallow rototilling (5-15 cm) including the application of dead mulch to potatoes under minimum tillage versus 2) conventional moldboard plough tillage (20-25 cm). The dead mulch applied to potatoes under minimum tillage was typically obtained from rye/pea or triticale/vetch cover crop mixtures that were chopped (< 10 cm) and applied at 10-15 t ha^−1^ on average with an adapted manure spreader. The C/N ratio of the mulch ranged between 20-25. Each tillage main plot (12 × 60 m^2^) was split into eight 6 × 15 m^2^ sub-plots of which four of the subplots received ∼5 t dry matter (ha yr)^−1^ of a high-quality yard waste compost that was applied manually. The remaining four sub-plots received potassium (K_2_SO_4_) and phosphorus (rock phosphate) fertilizer equivalent to the amounts of the composts. For details of the compost used between 2012 and 2015, see Schmidt et al. [21]. Composts used thereafter were of similar quality. In total, 32 plots were investigated (2 field experiments × 4 replicates × 2 tillage treatments × 2 fertilizer treatments).

**Table 1.**
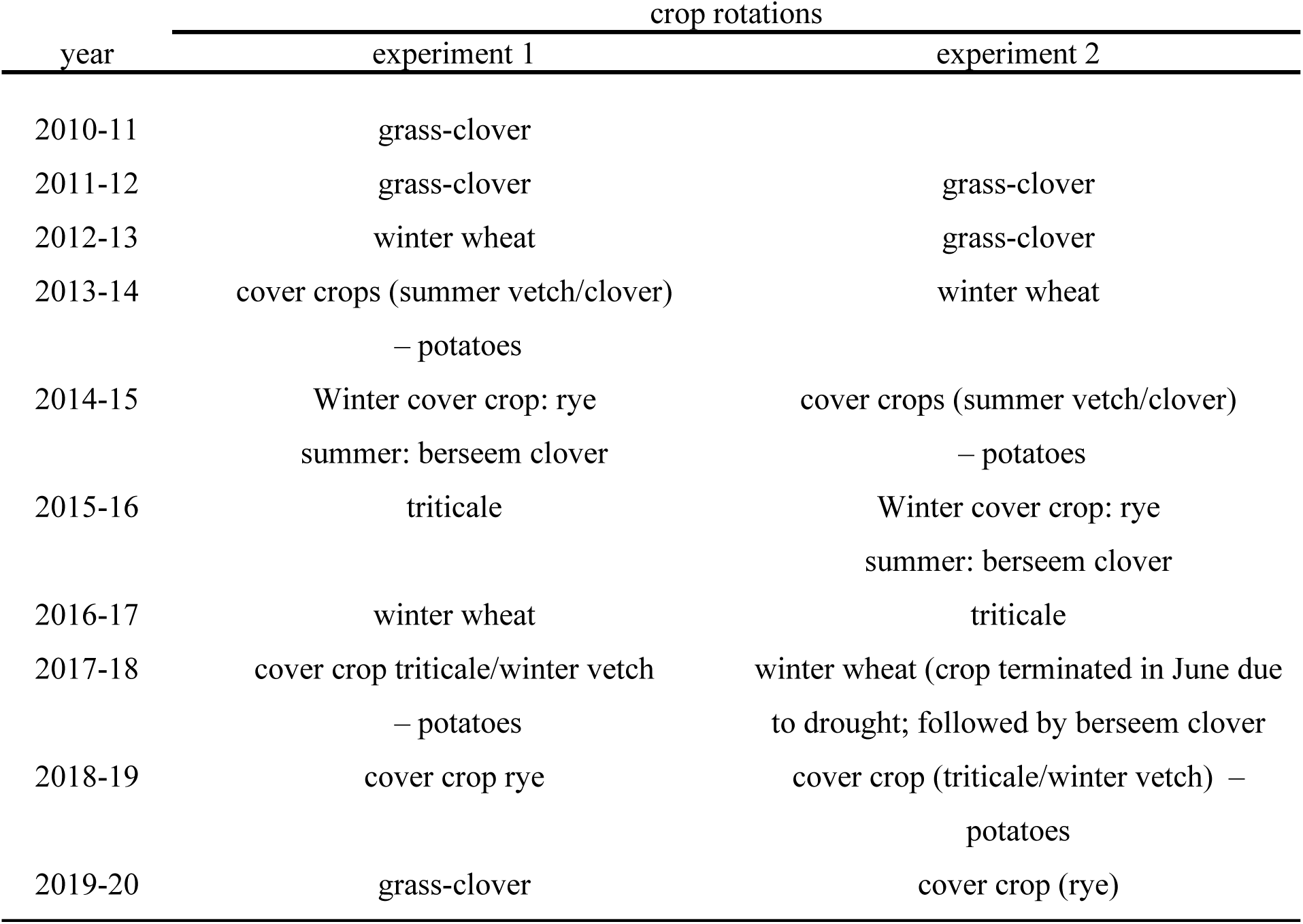
Sequence of crop rotations for the two field experiments including cash and cover crops (in brackets) from 2010 until 2020.

| year | crop rotations |  |
| --- | --- | --- |
|  | experiment 1 | experiment 2 |
| 2010-11 | grass-clover |  |
| 2011-12 | grass-clover | grass-clover |
| 2012-13 | winter wheat | grass-clover |
| 2013-14 | cover crops (summer vetch/clover)<br>– potatoes | winter wheat |
| 2014-15 | Winter cover crop: rye<br>summer: berseem clover | cover crops (summer vetch/clover)<br>– potatoes |
| 2015-16 | triticale | Winter cover crop: rye<br>summer: berseem clover |
| 2016-17 | winter wheat | triticale |
| 2017-18 | cover crop triticale/winter vetch<br>– potatoes | winter wheat (crop terminated in June due to drought; followed by berseem clover) |
| 2018-19 | cover crop rye | cover crop (triticale/winter vetch) –<br>potatoes |
| 2019-20 | grass-clover | cover crop (rye) |

Soil samples were collected after the first differential tillage (Experiment 1: Winter 2012/13, Experiment 2: Fall 2013), in September after the first potato crop (Experiment 1: 2014, Experiment 2: 2015), and in October 2019 (Experiment 1: during grass-clover crop, Experiment 2: after potatoes) (compare Table 1). Soil samples of the first two sampling dates of each experiment were taken from the top soil (0-15 cm) of each plot with a soil corer (2.4 cm diameter). In 2019, about 110 soil cores were taken randomly from each plot center (4.5 m × 10 m) with an Edelmann corer (8 cm diameter, 15 cm soil depth). Soils were sieved subsequently to 1 cm and stored in plastic bags at 4°C until processing.

### Soil nutrient analyses

Soil nutrients were analyzed by the Landesbetrieb Hessisches Landeslabor (LHL, http://www.lhl.hessen.de) two years after experimental setup, i.e. 2012 in experiment 1 and 2013 in experiment 2, and for both experiments in 2019. The former analysis was performed on pooled samples of each experimental field and pH and macronutrients were determined. In 2019, aliquots of 500 ml soil of each plot were analyzed for macro- and micronutrients, soil organic carbon (C_org_), total soil nitrogen (N_total_), pH, and salt (KCl) according to the “Verband Deutscher Landwirtschaftlicher Untersuchungs-und Forschungsanstalten” (VDLUFA) standards [22]: Book 1 A 4.1.3.2: direct assessment of C_org_ by burning at 550°C, Book 1 A 5.1.1: pH in soil-salt (CaCl) solution, Book 1 A 6.2.1.1: assessment of phosphorous and potassium in an acidic calcium-acetate-lactate-solution (CAL), Book 1 A 6.2.4.1: extraction of magnesium with CaCl_2_-solution and subsequent photometric detection, Book 1 A 10.1.1: calculation of KCl contents after assessment of electric conductivity, DIN EN ISO 17294-2:2017: determination of copper, zinc and boron in water via inductively-coupled plasma mass spectrometry, DIN EN ISO 11885:2009: determination of manganese and iron in water via inductively coupled plasma-optical emission spectrometry, and DIN EN 16168:2012: assessment of N_toal_ via dry burning.

### Biological assessments

Microbial biomass was determined two years after start of the experiments and directly after the first differential tillage (2012 and 2013), four years after start of the experiments (2014 and 2015), and 9 and 8 years after start of experiment 1 and 2 (2019), respectively. Soil samples were sieved to 2 mm and soil moisture was measured gravimetrically after drying at 105°C. After removing plant roots, soil microbial biomass was calculated via chloroform-fumigation extraction, following the instructions and equations of Vance et al. [23]. The resulting microbial biomass carbon (C_mic_) and nitrogen (N_mic_) values were divided by 0.45 and 0.54, respectively, which are the correction factors of the extractable microbial biomass in soils [24,25]. Extracts were stored at −15°C until organic C and total N in extracts were measured using an automatic C- and N-analyzer (Multi C/N, Fa Analytik Jena).

The microbial respiration as indicator for microbial activity, was determined from the sieved soils in 2019. Soils were moistened to 50% water-holding capacity for seven days prior to analysis. Two 70 g sub-samples of each soil were then filled into glass beakers placed in preserving jars that contained 20 ml water to prevent drying of the soils. Glass beakers with 15 ml of 0.5 mol NaOH were additionally placed in the jars. Six blinds without soil were used as controls. Jars were closed hermetically and incubated for seven days at 20°C. After incubation, glass beakers with NaOH were stored in vacuum desiccators filled with soda-lime to avoid evaporation of the CO_2_. The total CO_2_ concentration in the NaOH was assessed via HCl titration. For this, a solution containing 3 ml of the NaOH, 30 ml water, 3 ml 0.5 mol BaCl_2_, and two drops of phenolphthalein was stirred and titrated with 0.1 mol HCl until color change to rose. This back-titration will titrate the excessive NaOH. The soil respiration was calculated according to the formula:

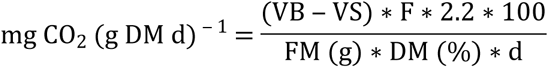

VB and VS are the volumes of HCl titrated to the blinds and samples, respectively, F is the dilution factor (3 ml aliquot of the 15 ml NaOH samples means F = 5), 2.2 corresponds to the amount of CO_2_ (mg) that refers to 1 ml of the titrated 0.1 mol HCl, FM and DM are the fresh matter (g) and dry matter (%) of the soil samples, respectively, and d is the incubation time (days) of the samples at 20°C.

For nematode analysis, 250 ml soil aliquots were processed with the Oostenbrink elutriator [26]. Nematodes collected on three mounted 45 µm sieves were washed into a beaker and transferred onto an Oostenbrink dish to get a clean sample. After incubation at room temperature for 48 hours, nematodes in the Oostenbrink dish were collected on a 20 µm sieve and transferred to a plastic tube and filled up to 30 ml with tap water. Nematode densities were counted from three times 1 ml aliquots at 40x magnification under a compound microscope. Final nematode densities were given as nematodes 100 ml soil^−1^. Nematodes were classified to the family level by morphological identification of 100 individuals sample^−1^ using the key of Bongers [27]. Nematode families were summarized into bacterivorous, fungivorous, herbivorous, and omnivorous/predatory feeding types by using the nematode indicator joint analysis web tool [28].

### Greenhouse study for fertility assessments

The soil fertility of each of the 32 soil samples (2 experiments × 4 replicates × 4 soils) was examined in a pea bioassay under greenhouse conditions at 21°C/18°C day/night temperatures. To check if difference in soil fertility might be associated with better plant performance under nematode pressure, the soil was artificially inoculated with the root lesion nematode *Pratylenchus penetrans*. For this, five sub-samples with 700 ml soil each were filled in 11×11×12 cm pots and the pots were organized as a randomized complete block (160 pots). Five surface-sterilized (70% ethanol for 5 minutes) and pre-germinated (2 days) pea seeds were planted per pot and reduced to three plants per pot after one week. At the day of plant reduction, mixed stages of *P. penetrans* (males, females, juveniles, and eggs) were inoculated in all soils at densities of 1000 nematodes and eggs 100 ml soil^−1^. The inoculation density was based on repeated pre-experiments with inoculation densities of 0, 500, 1000, 2000, and 3000 nematodes 100 ml soil^−1^ of the experiment 1 field. The pea biomass reduction was 11% and 12% in the pre-trial 1 and 2, respectively, after inoculation with 1000 *P. penetrans* 100 ml soil^−1^.

Nematode inoculum was obtained from 8-12 weeks old carrot discs cultures, that were chopped and extracted with Oostenbrink dishes [29]. The nematodes were stored at 7°C until used in the experiment.

Pots were watered to approximately 50% water-holding capacity every two to three days. The four tables with the pots were re-randomized each week to compensate for differences in illumination from neighboring cabins. Plants were harvested at BBCH 71 after a growth period of 80 days and aboveground dry matter (DM) after heating at 105°C for 24 h, the number of pods, and root fresh weight were determined. A root rot disease index (0-100) was calculated based on the assessment of external root lesions and lesion lengths were measured according to Šišić et al. [30] and Pflughöft [31]. *Pratylenchus penetrans* was extracted from pea roots via mist chambers [26]. For this, roots were cut in 1 cm pieces and placed on sieves that were placed on glass Petri dishes. Roots were kept moist for four weeks by spraying with water for 30 sec every 5 min. Once a week, nematodes settled on the ground of the Petri dish were transferred into PET bottles and stored at 4°C until the end of the extraction procedure. The final suspension was adjusted to 50 ml and 3 ml aliquots were taken to count the number of *P. penetrans*.

### Data processing and statistical analysis

Statistical analyses were performed with R version 3.6.0 [32], using the packages ‘nlme’ [33] for analyses via linear mixed models and ‘emmeans’ [34] for multiple comparison of treatments, back-transformed means, and standard errors. The package ‘car’ [35] was used to test the applied models for their variance homogeneity via Levene-tests. In case of violations of variance homogeneity, linear mixed effects models (lme) were adjusted with the weighting function ‘varIdent’ [36]. This function enables the model to use individual standard errors for each factor level (combination). If more than one factor showed inhomogeneous variances, the model with the lowest Akaike information criterion values was used based on log likelihood tests according to the description of Zuur et al. [36]. Fixed factors were tillage and fertilizer whilst random factors were experiments, field replicates, and tillage, each nested in the preceding factor. Thus, the formula used in R including an example of a weighting function with two factors that showed inhomogeneous variances was:

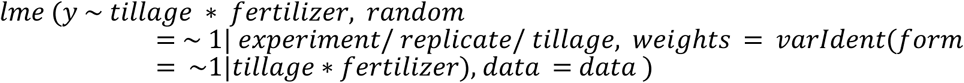

For the data analysis of the greenhouse experiment, the random term was extended to “random = ∼ 1|experiment/ replicate/ (greenhouse replicate/ tillage)”.

Spearman’s ρ rank correlations were used to study the relationship of chemical and biological indicators as well as their correlation with pea biomass production, root disease severity, and the number of *P. penetrans* in roots by using the ‘rcorr’ function of the R-package ‘Hmisc’ [37]. Results were visualized for each field experiment separately using the R function ‘corrplot’ of the ‘Hmisc’ package based on the *P* < 0.05 significance level.

## Results

Both field experiments were maintained according to the *ceteris paribus* principle. However, the severe drought in 2018 required some modifications in experiment 2. Due to the drought and also high weed infestation, the winter wheat was terminated two months earlier than usual (Table 1). The following summer cover crop (berseem clover) did not germinate because of the drought and therefore, a triticale/winter vetch cover crop was sown about two months after termination of the winter wheat.

Due to the time shift between the two field experiments (e.g. last application of dead mulch in experiment 1 and 2 dated back 16 and 4 months, respectively), we first added the experiment as fixed factor in the linear model. Hence, interactions of field experiment with tillage and fertilizer could be analyzed. In detail, experiments interacted with tillage regarding boron, KCl, C_mic_/C_org_, bacterivorous, and fungivorous nematodes (F_1,6_ > 8.1, *P <* 0.03, Table 2) but not with compost for any of the determined soil parameters. The interactions were, with the exception of fungivorous nematodes, expressed by a lower differentiation of minimum tillage from plough tillage in experiment 2 compared to experiment 1. As an example, boron concentrations were 51% and 14% higher under minimum tillage compared to plough tillage in experiment 1 and 2, respectively (data not shown). This justified the analysis of treatment effects across both field experiments with experiment as random factor.

**Table 2.** Means (± SE) of chemical and biological soil parameters in 2019, nine (experiment 1) and eight (experiment 2) years after start of the field experiments. Factors studied were plough and minimum tillage combined with either compost fertilization (< 5 t (ha a)^−1^) or mineral potassium and phosphorous application equivalent to contents in the respective composts. Results are averaged across both independent field experiments as only few experiment (Exp) by tillage (T) and no Exp by fertilizer (not shown) interactions occurred. Pea dry matter was obtained from a separate greenhouse experiment with the same soil used for analyses of the other parameters. Mean values for each soil parameter that do not share a common letter are significantly different (P < 0.05, df_tillage_ = 7, df_fertilizer between tillage_ = 7, df_fertilizer within tillage_ = 14) according to linear mixed effects models and estimated marginal means with Tukey corrections for multiple testing. ^1^Omnivorous (Om) and predatory (Pr) nematodes were pooled due to their overall low abundance.

| Soil parameter | unit | plough tillage |  | minimum tillage |  | P < F<br>(Exp x T) |
| --- | --- | --- | --- | --- | --- | --- |
|  |  | mineral | compost | mineral | compost |  |
| pH | | 6.56 ± 0.20 | 6.74 ± 0.20 | 6.70 ± 0.20 | 6.74 ± 0.19 | n.s. |
| P<sub>2</sub>O<sub>5</sub> | mg (kg soil)<sup>-1</sup> | 71 ± 28a | 103 ± 29ab | 125 ± 28b | 133 ± 29b | n.s. |
| K<sub>2</sub>O | mg (kg soil)<sup>-1</sup> | 124 ± 9.4a | 136 ± 11.2a | 319 ± 9.1b | 324 ± 23.5b | n.s. |
| MgO | mg (kg soil)<sup>-1</sup> | 166 ± 11.9a | 174 ± 12.1ab | 172 ± 11.8a | 189 ± 12.4b | n.s. |
| Cu | mg (kg soil)<sup>-1</sup> | 3.29 ± 0.22b | 2.99 ± 0.22a | 3.07 ± 0.23ab | 3.15 ± 0.26ab | n.s. |
| Zn | mg (kg soil)<sup>-1</sup> | 5.35 ± 0.25a | 6.5 ± 0.47b | 6.94 ± 0.25b | 7.15 ± 0.66ab | n.s. |
| Mn | mg (kg soil)<sup>-1</sup> | 374 ± 34 | 354 ± 34 | 394 ± 36 | 357 ± 34 | n.s. |
| B | mg (kg soil)<sup>-1</sup> | 0.65 ± 0.11a | 0.71 ± 0.11a | 0.89 ± 0.11b | 0.91 ± 0.11b | 0.001 |
| Fe | mg (kg soil)<sup>-1</sup> | 118 ± 13 | 117 ± 13 | 121 ± 13 | 120 ± 13 | n.s. |
| Salt (KCl) | mg l<sup>-1</sup> | 486 ± 49a | 582 ± 67abc | 700 ± 49b | 762 ± 50c | 0.03 |
| C<sub>org</sub> | % | 1.27 ± 0.06a | 1.52 ± 0.07b | 1.79 ± 0.06c | 1.95 ± 0.07d | n.s. |
| N<sub>total</sub> | % | 0.15 ± 0.007a | 0.18 ± 0.008b | 0.2 ± 0.006c | 0.21 ± 0.008c | n.s. |
| Microbial respiration (MR) | $\mu\text{g CO}_2$ (g dry soil d)<sup>-1</sup> | 33.9 ± 1.3a | 41.6 ± 1.2b | 66.7 ± 4.4c | 73.7 ± 3.7c | n.s. |
| C<sub>mic</sub>/N<sub>mic</sub> | | 5.62 ± 0.27 | 5.52 ± 0.35 | 4.76 ± 0.12 | 4.84 ± 0.19 | n.s. |
| C<sub>mic</sub>/C<sub>org</sub> | % | 2.47 ± 0.19ab | 2.36 ± 0.19a | 2.77 ± 0.22ab | 2.7 ± 0.2b | 0.02 |
| MR/C<sub>mic</sub> | % | 11.1 ± 1.27a | 11.7 ± 1.27ab | 13.5 ± 1.27ab | 14.0 ± 1.27b | n.s. |
| N<sub>mic</sub>/N<sub>total</sub> | % | 3.68 ± 0.31a | 3.86 ± 0.31a | 5.27 ± 0.31b | 5.18 ± 0.31b | n.s. |
| Bacterivorous nematodes | individuals 100 ml soil<sup>-1</sup> | 546 ± 471a | 662 ± 469a | 1263 ± 478b | 1295 ± 474b | 0.03 |
| Fungivorous nematodes | individuals 100 ml soil<sup>-1</sup> | 326 ± 69 | 313 ± 69 | 417 ± 69 | 335 ± 69 | 0.008 |
| Herbivorous nematodes | individuals 100 ml soil<sup>-1</sup> | 568 ± 149 | 675 ± 149 | 889 ± 149 | 924 ± 149 | n.s. |
| Om + Pr nematodes<sup>1</sup> | individuals 100 ml soil<sup>-1</sup> | 72 ± 21 | 118 ± 21 | 96 ± 36 | 155 ± 36 | n.s. |

### Effects of tillage system and fertilizer application on chemical and biological soil properties

Minimum tillage and in part also compost application increased the amounts of most macronutrients in soil. Initial contents of P_2_O_5_, K_2_O, MgO, and C_org_ in the top 25 cm soil two years after start of the field experiments (2012/2013) were on average 105, 140, 145 (all in mg (kg soil)^−1^), and 1.2%, respectively (data not shown). In 2019, these values were lower or similar under plough tillage with mineral fertilization (Table 2). In comparison, values were slightly higher under plough tillage with compost fertilization, in particular, C_org_ was increased by 20% (Table 2). In contrast, minimum tillage with compost or mineral fertilization increased P_2_O_5_, K_2_O, C_org_ by 23%, 129%, and 57% compared to initial values, respectively. This translated to 48%, 147%, and 34%, higher P_2_O_5_, K_2_O, and C_org_ values, respectively, under minimum tillage compared to plough tillage in 2019, regardless if mineral or compost fertilization was applied (Table 2). Moreover, N_total_ was 25% higher under minimum tillage than plough tillage (Table 2).

In the present study, pH was considerably higher in all treatments in 2019 than when first measured in 2012 and 2013 (6.1, not shown). Differences in pH between treatments were not observed.

The salt content varied between 486 (plough tillage without compost) and 762 (minimum tillage with compost) mg KCl l^−1^, which translates to electrical conductivities (EC) of 0.9 and 1.4 mS cm^−1^, respectively.

Soil micronutrients varied in a distinct pattern among treatments. While copper concentrations were highest under plough tillage with mineral fertilization, manganese and iron concentrations were similar across all treatments (Table 2). In contrast, zinc and boron concentrations were significantly higher under minimum than under plough tillage, irrespective of the fertilization strategy.

In general, soil biological properties were enhanced by minimum tillage compared to the plough tillage systems. For C_mic_, the differences between minimum and plough tillage as well as in part between the compost and mineral fertilization increased over time (Fig 2). This is reflected by significant interactions of sampling date (year) and tillage in both experiments (F_2,30_ > 7.7, *P* ≤ 0.002). The *status quo* analysis was taken after the first differential tillage and compost application (2012, 2013), two years after the start of the experiment. Initial C_mic_ values in experiment 1 and 2 were 60% and 27% higher under minimum tillage with mineral fertilizer than under plough tillage with mineral fertilizer. However, those differences were not statistically significant due to large standard errors (Fig 2).

**Fig 2.**
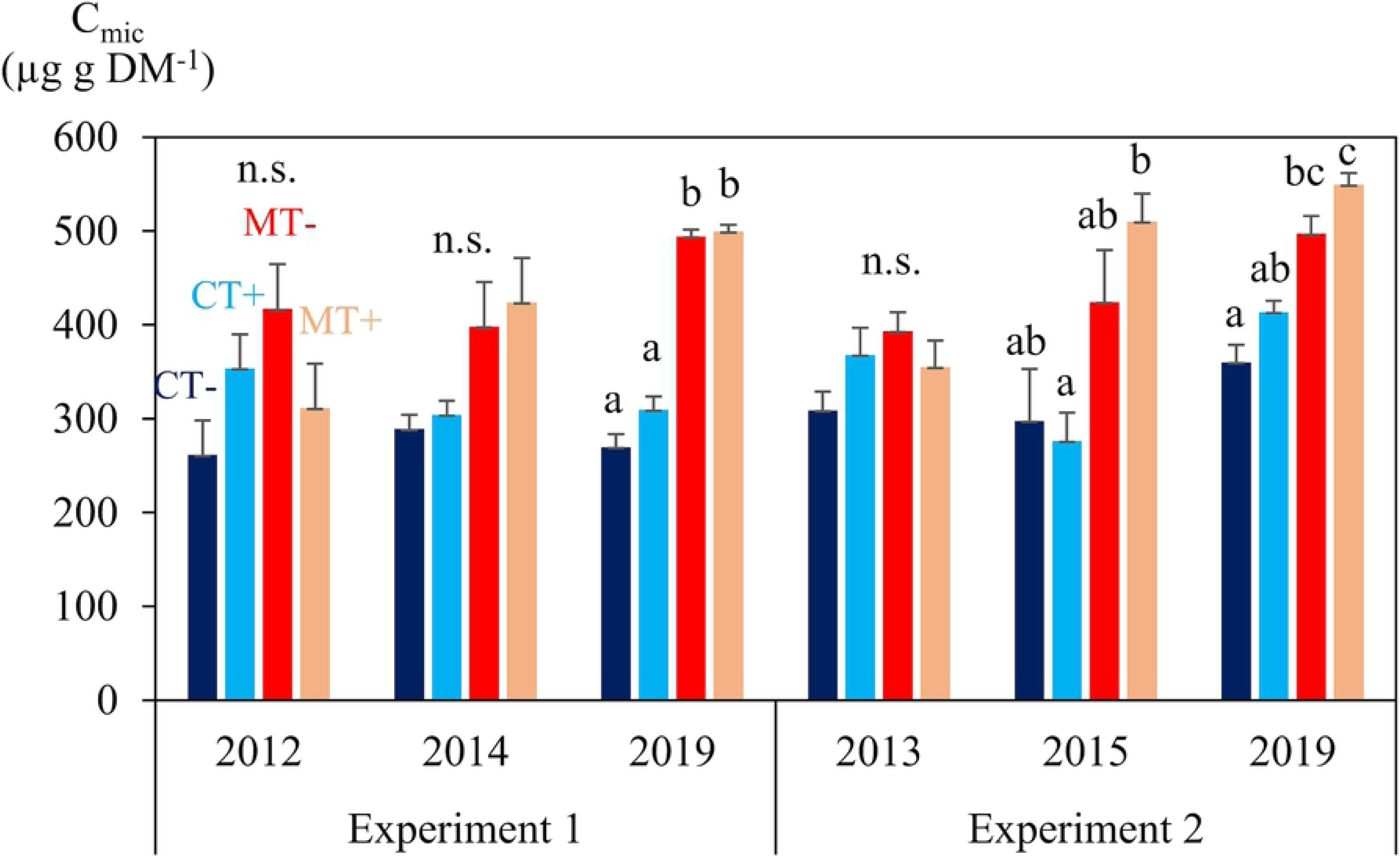
Mean (± SE) top soil (15 cm) microbial biomass carbon (C_mic_) in µg g soil dry matter^−1^ affected by plough (CT, blue bars) or minimum tillage (MT, red bars) each combined with either yard waste compost application (+, light colored bars) or mineral potassium and phosphorous application equivalent to contents in the composts (-, dark colored bars). C_mic_ was determined two, four years, and eight/nine years after start of the experiments (experiment 1: 2010; experiment 2: 2011). Mean values of the respective treatments that do not share a common letter for each year and experiment are significantly different (P <0.05, df_tillage_= 3, df_fertilizer_= 30) according to linear mixed effects models and estimated marginal means with Tukey correction for multiple testing.

Four years after the start of the experiment when potatoes had been grown for the first time with mulch application, C_mic_ was already 39% and 62% higher under minimum tillage than under plough tillage in experiment 1 (2014) and 2 (2015), respectively (Fig 2). At that time, compost application had increased C_mic_ consistently (6-20%) in comparison to mineral fertilization in both experiments under minimum tillage.

In 2019, C_mic_ values were 72% and 35% higher under minimum compared to plough tillage in experiment 1 and 2, respectively. The C_mic_ was 15% higher under plough tillage with compost compared to plough tillage with mineral fertilization (Fig 2). Compost effects under minimum tillage were less pronounced in 2019 than in 2014 and 2015. Similar effects of tillage were observed for the microbial respiration and number of free-living nematodes in both years that were on average 86% and 64% higher, respectively, under minimum than plough tillage (Table 2). In particular, the number of bacterivorous nematodes was three-fold and two-fold higher under minimum tillage compared to plough tillage in experiment 1 and experiment 2, respectively, which also explains the significance of the experiment by tillage interaction (Table 2). The absolute number of herbivorous as well as omnivorous/predatory nematodes was 31% to 46% higher under minimum tillage compared to plough tillage. Fungivorous nematodes showed no statistical differences among treatments. However, total numbers of fungivorous nematodes showed a different pattern in both years. In the first year (experiment 1) the total number of fungivorous nematodes was higher under minimum tillage (425 nematodes 100 ml soil^−1^) compared to plough tillage (173 nematodes 100 ml soil^−1^), whereas in the second year it was the opposite with higher numbers under plough tillage (466 nematodes 100 ml soil^−1^) compared to minimum tillage (327 nematodes 100 ml soil^−1^).

### Effects of tillage system and fertilizer application on pea performance under greenhouse conditions

Overall, pea yield was similar in both experiments. For example, pea aboveground dry weight was 50% and 39% higher under minimum tillage compared to plough tillage in experiment 1 and 2, respectively. Although this effect was less clear for the pea root fresh weight, the highest root weights were generally recorded under minimum tillage (Table 3). Similarly, the number of pods produced per pot was always higher under minimum tillage than under plough tillage.

**Table 3.**
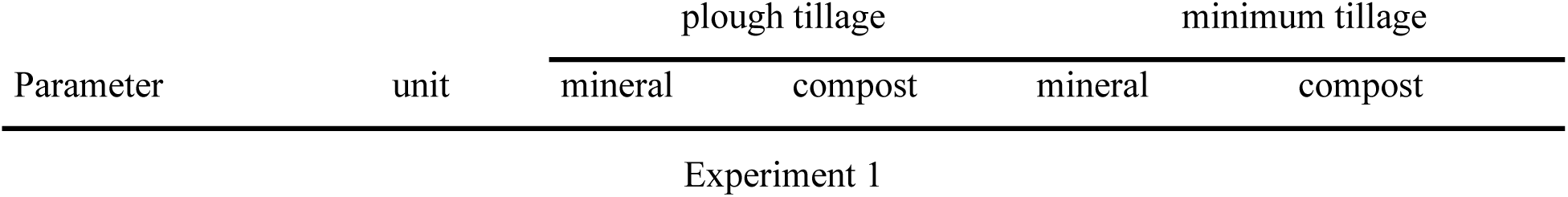
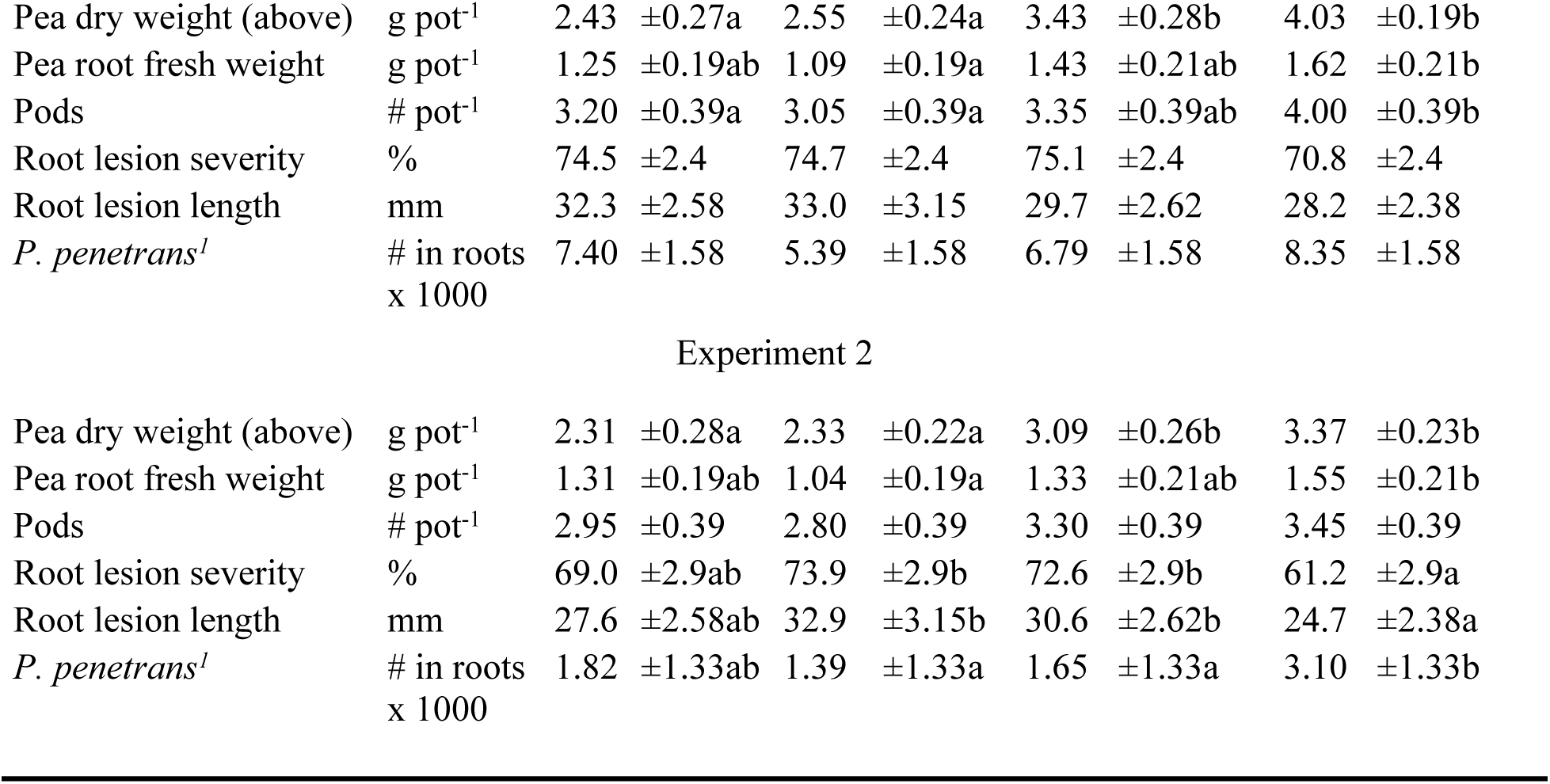
Estimated marginal means (± SE) of pea production parameters in both field experiments assessed in a greenhouse experiment. Factors studied were plough and minimum tillage combined with either compost fertilization (< 5 t (ha a)^−1^) or mineral potassium and phosphorous application equivalent to contents in the respective composts. Mean values for each parameter that do not share a common letter are significantly different (P < 0.05, df_tillage_ = 38, df_fertilizer between tillage_ = 38, df_fertilizer within tillage_ = 76) according to linear mixed effects models and estimated marginal means with Tukey corrections for multiple testing. ^1^Soils were inoculated with 7000 mixed stages + eggs of *P. penetrans* pot^−1^.

| Parameter | unit | plough tillage |  | minimum tillage |  |
| --- | --- | --- | --- | --- | --- |
|  |  | mineral | compost | mineral | compost |
| Experiment 1 |  |  |  |  |  |
| Pea dry weight (above) | g pot<sup>-1</sup> | 2.43 ±0.27a | 2.55 ±0.24a | 3.43 ±0.28b | 4.03 ±0.19b |
| Pea root fresh weight | g pot<sup>-1</sup> | 1.25 ±0.19ab | 1.09 ±0.19a | 1.43 ±0.21ab | 1.62 ±0.21b |
| Pods | # pot<sup>-1</sup> | 3.20 ±0.39a | 3.05 ±0.39a | 3.35 ±0.39ab | 4.00 ±0.39b |
| Root lesion severity | % | 74.5 ±2.4 | 74.7 ±2.4 | 75.1 ±2.4 | 70.8 ±2.4 |
| Root lesion length | mm | 32.3 ±2.58 | 33.0 ±3.15 | 29.7 ±2.62 | 28.2 ±2.38 |
| *P. penetrans*<sup>l</sup> | # in roots<br>x 1000 | 7.40 ±1.58 | 5.39 ±1.58 | 6.79 ±1.58 | 8.35 ±1.58 |
| Experiment 2 |  |  |  |  |  |
| Pea dry weight (above) | g pot<sup>-1</sup> | 2.31 ±0.28a | 2.33 ±0.22a | 3.09 ±0.26b | 3.37 ±0.23b |
| Pea root fresh weight | g pot<sup>-1</sup> | 1.31 ±0.19ab | 1.04 ±0.19a | 1.33 ±0.21ab | 1.55 ±0.21b |
| Pods | # pot<sup>-1</sup> | 2.95 ±0.39 | 2.80 ±0.39 | 3.30 ±0.39 | 3.45 ±0.39 |
| Root lesion severity | % | 69.0 ±2.9ab | 73.9 ±2.9b | 72.6 ±2.9b | 61.2 ±2.9a |
| Root lesion length | mm | 27.6 ±2.58ab | 32.9 ±3.15b | 30.6 ±2.62b | 24.7 ±2.38a |
| *P. penetrans*<sup>l</sup> | # in roots<br>x 1000 | 1.82 ±1.33ab | 1.39 ±1.33a | 1.65 ±1.33a | 3.10 ±1.33b |

Root lesion severity and root lesion length were about 5% higher in experiment 1 compared to experiment 2. In general, both diseases parameters were similar for the two plough tillage systems and the minimum tillage system that had received mineral fertilizer. However, when peas were grown in soil collected from the minimum tillage system that was fertilized with compost root lesion severity and root lesion length were reduced. This effect was only significant in experiment 2, though.

The number of *P. penetrans* in roots pot^−1^ was 6,983 and 1,990 in experiment 1 and 2, respectively. Hence, the final population density divided by the inoculation density was 1 and 0.29, respectively. In both experiments, the number of *P. penetrans* was lowest under plough tillage fertilized with compost and highest under minimum tillage fertilized with compost.

### Biological soil components as indicators of soil fertility

In our study, soil fertility was measured as the potential of the soils for pea biomass production in a greenhouse bioassay after artificial soil inoculation with lesion nematodes (Table 3).

The growth substrate of pea in the greenhouse and the soil used for the analysis of the soil chemical and biological parameters shown in Table 2 were derived from the same composite samples. This allows to directly link these with data from the greenhouse experiment that were used as ecosystem services (Fig 3). Thus, a number of standard biological “black box” indicators of soil quality, such as C_mic_, N_mic_, microbial respiration, and C_org_ were positively correlated with the pea biomass production in soils of both field experiments (Fig 3). The severity of root lesions further affected pea dry matter in both experiments (ρ < −0.51, *P* < 0.46, Fig 3).

**Fig 3.**
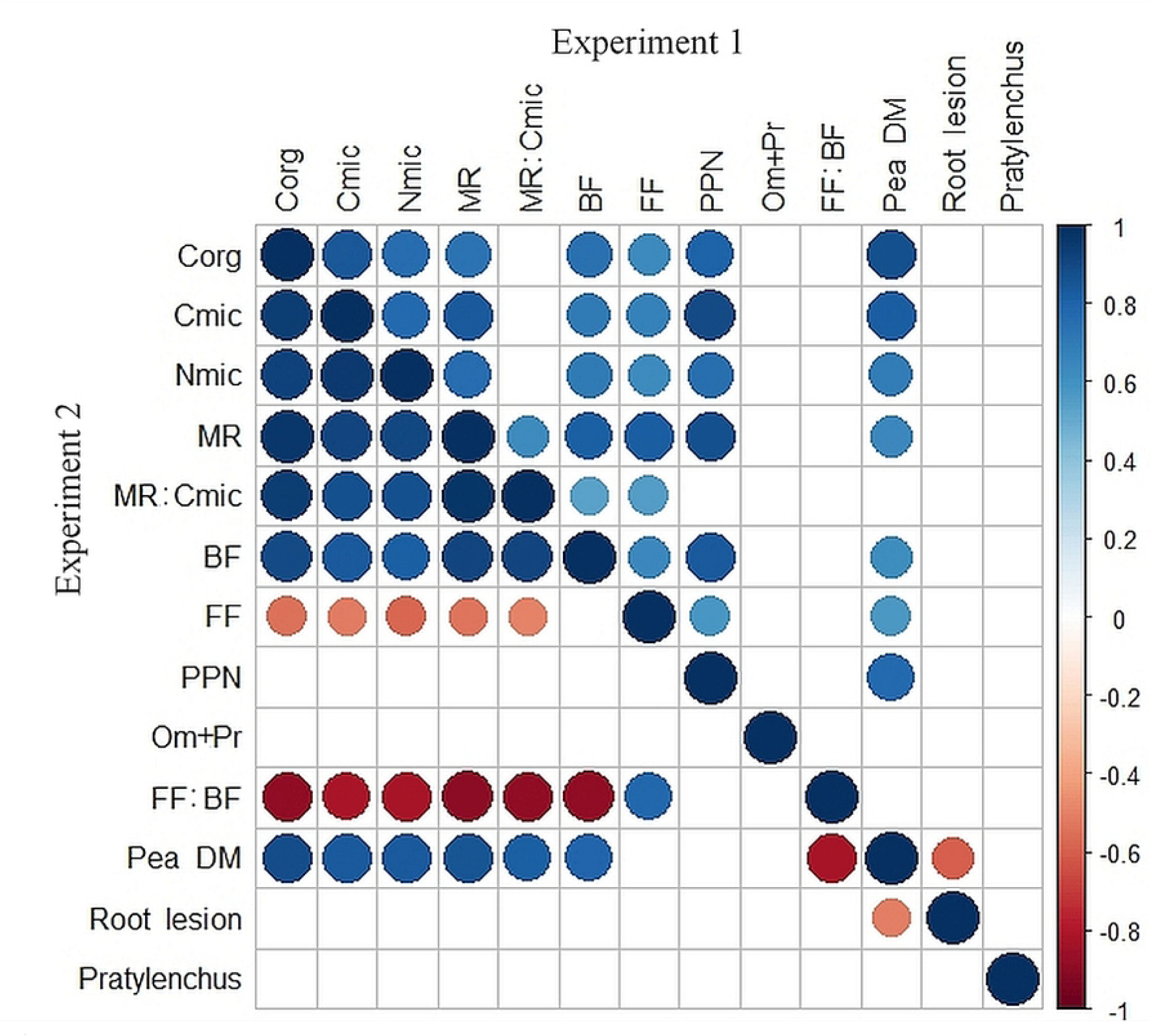
Spearman’s ρ correlation coefficients of important biological soil indicators (including C_org_) independent of applied treatments. Correlations within experiment 1 and 2 are shown in the upper and lower diagonal, respectively. Positive correlations are displayed in blue and negative correlations in red color. Color intensity and the size of the circle are proportional to the correlation coefficients. In the right side of the correlogram, the legend color shows the correlation coefficients and the corresponding colors. Correlations with P-values > 0.05 (n= 16) are considered insignificant and were left blank. Abbreviations: Corg, total organic C; Cmic, microbial biomass C; Nmic, microbial biomass N; MR, microbial respiration; BF, bacterivorous nematodes; FF, fungivorous nematodes; PPN, herbivorous nematodes; Om+Pr, omnivorous and predator nematodes; Pea DM, pea dry matter; Root lesion, pea root rot; *Pratylenchus, Pratylenchus* in pea roots. The latter three indicators were assessed in a separate greenhouse experiment.

Besides these indicators, we also used the abundance of free-living nematodes that include free-living herbivorous, bacterivorous, fungivorous and omnivorous/predatory nematodes. In both experiments, the abundance of bacterivorous nematodes was highly correlated with pea dry matter (ρ > 0.61, *P <* 0.012, Fig 3) as well as with C_mic_ and microbial respiration (ρ > 0.7, *P <* 0.003, Fig 3). The pooled abundance of omnivorous/predatory nematodes as well as the number of *P. penetrans* in roots pot^−1^ were unaffected by any of the applied parameters. Herbivorous nematodes were positively correlated with pea dry matter in experiment 1 (ρ > 0.78, *P <* 0.001, Fig 3) but not in experiment 2 (ρ > 0.23, *P > 0*.05, Fig 3). In experiment 2, fungivorous nematodes correlated negatively with microbial biomass, respiration and C_org_ while the fungivorous:bacterivorous ratio correlated with reduced pea dry matter yields (ρ > −0.83, *P <* 0.001). A minor negative correlation (ρ > −0.3, *P >* 0.17) was observed among root lesion severity at the stem base and on roots of pea and the number of *P. penetrans* in roots, however, this effect was not statistically significant in any of the experiments.

## Discussion

In general, nutrient availability is one of the major yield limiting factors in organic agriculture due to the restrictions on the use of soluble mineral fertilizers. Under these conditions, nutrient deficiency can be even worse when accompanied by minimum tillage [7]. The reason is that aboveground soil cover delays soil warming and thus, nutrient mineralization, which can severely affect early seedling development [2]. The minimum tillage system presented here, combining cover crops and dead mulch applications clearly showed the potential to overcome such nutrient deficiencies. This is the result of greater soil fertility in the top layer compared to the initial levels as well as to the plough-based system. Interestingly, potato yields correlated positively with the numbers of free-living nematodes determined in 2019 [6]. In addition to greater soil fertility in the top layer, mulching is an important measure to conserve water further leading to yield increases, a phenomenon we have consistently observed with mulched potatoes in the years 2015 and 2016 [38] and also in 2018 and 2019 (Junge, Finckh, et al., unpublished data). The improvement in soil fertility was similar or even greater compared to a long-term organic study from Frick, Switzerland, where phosphorus and potassium were 72% and 40% higher in the top 10 cm soil following six years of reduced tillage (15 cm deep two-layer ploughing) compared to 30 cm deep ploughing [39]. In the same study, C_org_ increased from 2.2% to 2.6% under reduced tillage, whereas it remained at 2.1% at deep ploughing. After 15 years of minimum tillage in the same experiment, C_org_ contents were increased from 2.2% to 2.9% [5], which is similar to the increase in our study after 8 and 9 years of minimum tillage.

With respect to the similarity of the pH levels across tillage and fertilizer applications, it could not be determined if there were no differences per se or if liming of the two field experiments in the month before sampling camouflaged tillage or fertilizer effects on pH. However, in a similar experiment in Austria, where shallow conservation tillage was compared to 25 cm deep ploughing, also no significant differences in pH were found [40]. Likewise, neither tillage nor slurry or composted manure had any effect on soil pH in the above described long-term study from Frick, Switzerland [5].

The observed higher salt values under minimum tillage should be seen with caution as elevated salt concentrations can cause yield depression of salt intolerant plants, such as described for Phaseolus bean and carrots, where threshold levels of about 1 mS cm^−1^ are given [41]. Comparing the fertilization systems in this study, higher salt contents were generally found for organic fertilization than for mineral fertilization. This is not surprising as the composts used had high salt contents (EC: 5-10 mS cm^−1^) [21].

The effect of the tillage system on micronutrients has rarely been described. However, it appears that this generally follows the same pattern as observed for macronutrients, i.e. an enrichment under minimum/zero tillage compared to intensive tillage [42]. Unfortunately, nothing is known about the relevance of those differences in micronutrient concentrations for plant performance. It can only be speculated that the higher micronutrient levels as well as macronutrient levels observed in this study, may have contributed to the greater biomass production in the pea bioassay under both minimum tillage systems compared to plough tillage (Table 2). For example, boron (and calcium, but this has not been assessed here) is known to be important for N_2_ fixation in legumes and can massively enhance pea aboveground biomass production [43]. However, if boron contributed to pea growth in this study remains unsolved.

With respect to biological soil properties, our results confirm other studies that showed that C_mic_ and microbial activity under minimum tillage are generally higher than under plough tillage [5,39,44]. The fact that this was shown in two independent field experiments under two different crops and also for the abundance of bacterivorous nematodes in the present study indicates the robustness of such measurements. Although soil biological properties can vary greatly across the season, Kandeler and Böhm [44] demonstrated that C_mic_ can already be used as a reliable indicator for changes in soil biology four years after the transition to minimum tillage. This is in line with the findings of our study, where the first clear differentiations between tillage systems occurred two years after differential tillage and four years after minimum tillage (Fig 2, year 2014 and 2015). Besides, we found higher microbial quotients (C_mic_/C_org_) under minimum than under plough tillage. These likely indicate a higher C input as well as a higher C quality for C_mic_ production [45].

Contrasting to our expectations, the microbial quotient (MR/C_mic_), which can be used as indicator of changes in organic matter availability [46], was 21% higher under minimum tillage compared to plough tillage. This suggests that minimum tillage in combination with dead mulch application induced a lower efficiency in C use compared to the plough system. C_org_ availability was also found to be one of the main drivers of microbial quotients in a study that compared 30 different textured soils with varying C_org_ contents [47]. Haynes [46] noted that bacteria are less efficient in utilizing C sources than fungi, which may further explain the higher metabolic quotient under minimum tillage compared to plough tillage in our study. The application of dead mulch with a low to moderate C/N ratio (∼25) was likely responsible for a domination of the break down of crop residues by bacteria. This is supported by a 3-fold greater bacterivorous:fungivorous nematode ratio under minimum tillage 4 months after dead mulch application (experiment 2) compared to 16 months after dead mulch application (experiment 1) (data not shown). Specifically, the number of bacterivorous r-strategist Rhabditidae and Panagrolaimidae were enhanced under minimum tillage by 44% in experiment 1 and 88% in experiment 2 when compared to plough tillage (unpublished data). In contrast to experiment 1, the recent disturbances in the minimum tillage system in experiment 2 due to potato production, i.e. tillage for planting, fertilization with nitrogen rich mulch and harvesting tillage likely fostered enrichment opportunists, i.e. bacterivorous nematodes of Rhabditidae and Panagrolaimidae [18,19]. This is confirmed by a study from Canada, where conservation tillage resulted in 17.5% to 119% higher numbers of bacterivorous nematodes (Rhabditidae and Diplogasteridae) compared to conventional tillage after potato cropping [48]. The lower C_mic_/N_mic_ ratios under minimum compared to plough tillage (Table 2) also suggest a dominance of bacteria over fungi. This is in contrast to lower bacteria:fungi ratios observed under minimum tillage compared to plough tillage systems under similar climatic conditions [49]. The fact that in the study of Kuntz et al. [49] the assessment had been performed eight months after the last tillage operation could have led to a general succession from bacterial to fungal break down of residues. According to Bongers and Bongers [18] the number of Rhabditidae and Panagrolaimidae in soil will decline with decreasing nutrient availability and microbial activity, while the general opportunistic Cephalobidae will become dominant. This occurred in the minimum tillage system in experiment 1, 16 months after the last mulch application. Further succession under minimum tillage will likely result in fungal breakdown of crop residues and thus in an increase of opportunistic fungivorous nematode families (Aphelenchoididae, Aphelenchidae) [18,20]. A similar situation likely occurred in the plough tillage system without mulch fertilization in experiment 1, where the number of fungivorous nematodes was twice as high as bacterivorous nematodes (234 nematodes 100 ml soil^−1^, unpublished). The greater relative abundance of fungivorous as well as omnivorous/predatory nematodes under plough tillage compared to minimum tillage was also observed in a study that compared a no-till with a plough system in a maize-soy bean rotation [50].

The massively enhanced pea biomass production under minimum tillage in the greenhouse assay could be expected based on the generally higher macro- and micronutrient levels compared to the plough-based system. The strongly positive effects of compost fertilization under minimum tillage could neither be related to differences in nutrient composition nor to biological properties. Suppression of *P. penetrans* did not play a role in this context as its numbers were generally highest under minimum tillage with compost. It is likely that the nematode benefitted from higher root masses and plant nourishment in this treatment. At C_org_ contents of <1.4%, the application of 4 t ha^−1^ of composted pig manure reduced *P. penetrans* by on average 87% in two pot experiments cropped with sugar beets [51]. Conditions were similar in the plough system in our study but the reduction of *P. penetrans* in the compost treated soil was only 25%. In contrast, the root lesion severity was only reduced under minimum tillage with compost. These lesions were likely caused by soil inhabiting and pea pathogenic fungi that were commonly found in soils of the study site [14,30]. The negative correlation of root lesion severity and pea biomass production in this study suggests that soil health under minimum tillage with compost fertilization is overall increased. This was also directly translated into a greater soil fertility in terms of pea biomass production. As pointed out above, under field conditions when there is a lack of rainfall, effects of the mulch and increased soil organic matter contents on water retention may lead to further advantages in practice.

Especially bacterivorous and fungivorous nematodes contribute to soil fertility as decomposers that release nitrogen to the soil [52,53]. Both feeding types accounted for 50% and 65% of the total nematode community in experiment 1 and 2, respectively, which highlighted their importance for nutrient turnover and thus pea production in our study. Across the globe, organic carbon that is commonly used as indicator of soil fertility was found as one of the main drivers of nematode abundance [15,54]. This was also observed for bacterivorous nematodes in both field experiments in our study. However, the fungivorous:bacterivorous nematode ratio was negatively correlated with pea dry matter yields in experiment 2, which emphasizes that bacterivorous nematodes were more reliable indicators of soil fertility than fungivorous nematodes. This is further expressed by the strong positive correlations of bacterivorous nematodes and pea biomass production in both field experiments (Fig 3). Also, the strong correlations of bacterivorous nematodes with microbial biomass and microbial respiration in both field experiments highlights their usefulness as indicators of a microbially active soil. The fact that predominantly bacterivorous nematodes are categorized into the group of enrichment opportunists [18,55] further strengthen our hypothesis that these are important indicators of soil fertility.

## Conclusion

We conclude that minimum tillage accompanied by regular mulch applications in a system with frequent and diverse use of cover crops in the rotation provides a promising management strategy for sustainable organic crop production. Soil fertility can be improved when converting to minimum tillage, even under organic conditions, if the system is properly adapted. Clearly, organic inputs are crucial for success. Soil fertility could be related to a number of chemical and biological soil indicators that, in turn, were positively correlated with pea biomass. This was achieved through a complete revision and adaptation of the organic farming system when converting to minimum tillage. Such a process is comparable to the conversion from conventional to organic farming. Considering the still increasing C_mic_ under minimum tillage and its strong correlation with pea biomass production, further soil fertility improvements can be expected if the system is maintained further. The results suggest an accumulation of an increasing pool of active organic matter, i.e. the pool of soil fauna excluding their resting stages and dead organisms, under minimum tillage that not only explains the higher microbial respiration in soil but also the higher pea biomass production compared to the plough tillage system. We therefore hypothesize that this particular minimum tillage system based on maximum use of cover crops and additional transferred mulch to potatoes fosters microbial nutrient turnover from the labile C pool and thus, improves plant nutrition and plant growth.

In general, the use of compost as organic fertilizer should be preferred over mineral fertilization as together with compost relevant micronutrients are supplied to the system. However, compost improved only few chemical and biological soil properties in the plough and almost none in the minimum tillage systems. The latter was quite surprising as compost application to the minimum tillage system resulted in the highest pea biomass under controlled conditions. Indeed, compost reduced the disease severity of root and stem root of the peas and, in soils with low organic carbon contents, likely improved the resilience towards the resident pest *P. penetrans*. Additional physical, chemical, and biological soil properties besides those investigated in our study may also play a role and need to be investigated to fully understand the management-soil-plant interactions studied here. In addition, the overall resilience of the management systems towards important biotic (fungal pathogens) and abiotic (drought, heat) stressors need to be investigated in order to fully understand the ecosystem services provided by the system.

The simple assessment of free-living nematodes and their classification into feeding types provides a useful indicator for soil fertility. Thus, the bacterivorous nematodes were equivalent indicators of soil fertility than other typically used parameters such as C_org_, C_mic_, microbial respiration, and macronutrients. Of course, such analyses could be considerably strengthened by more detailed nematological investigations. The low laboratory equipment cost for simple free-living nematode assessments, e.g. nematodes can easily be obtained from Oostenbrink dishes, Baermann funnels, or Cobb’s sieves and counted and classified into feeding types under a microscope at 40x magnification [26], is an additional advantage of using nematodes as bioindicators. Nematode specific indices, such as maturity, channel, and structure indices as well as metabolic footprints will provide more details about the faunal composition that influences the fertility and resilience status of a soil [19,52,56]. The nematodes’ key positions in the soil food web will thus allow to track the different carbon pathways in the soil without the use of expensive and specialized equipment, such as needed for phospholipid fatty acid, chloroform fumigation, ergosterol, and other extractions. However, detailed taxonomic knowledge is required to identify the free-living nematodes to the family and genus levels. In addition, identification to the species level by using molecular methods will give a detailed overview of the contribution of single species to a certain ecosystem service, such as plant production, disease suppression or resilience. In accordance with many other studies, our results clearly demonstrate that nematodes harbor a great potential for characterization of management effects.

## Acknowledgements

The authors would like to thank Matthias von Ahn and Leonard Theisgen for their excellent assistance in nematode extraction and count, as well as Elsa Zwicker and Keo Sasha Rigorth for their excellent support in microbial biomass and activity assessments. Furthermore, the authors would like to thank Stephan Junge for provision of potato yield data and field pictures.

